# Harnessing historical collections to reconstruct the invasion history of *Microstegium vimineum* (Poaceae)

**DOI:** 10.64898/2026.09.29.755326

**Authors:** Fernando Hernández, Cameron W. Corbett, Lauren E. Kosslow, Cynthia D. Huebner, Chih-Hui Chen, Brandon T. Hendrickson, Bo Li, Pan Li, Paul M. Peterson, Kenji Suetsugu, Wen-Bin Yu, Nicholas J. Kooyers, Craig F. Barrett

## Abstract

- The evolutionary processes underlying successful invasions are often difficult to disentangle. Historical collections provide a temporal perspective, offering great potential to reconstruct invasion histories and identify the processes driving invasions.
- Japanese stiltgrass (*Microstegium vimineum*) is an invasive C_4_ grass introduced to eastern North America over a century ago and now established in 31 USA states. To reconstruct its invasion history, we leveraged low-coverage whole-genome sequencing of 142 herbarium and 330 contemporary samples spanning the temporal and spatial extent of the North American invasion and the native range in Asia.
- We identified three independent introductions in the USA: an awnless form in the South and two independent introductions of awned forms in the North, with limited admixture among introductions across space and time. In both regions, we observed high genetic diversity in the earliest available collections despite demographic bottlenecks and identified several putative selective sweeps in or near flowering-time genes, suggesting rapid evolution in the introduced range.
- Our analyses support separate invasions in the USA, likely facilitated by standing genetic variation and selection, but provide little support for admixture playing a major role. Our study highlights the power of historical collections to uncover the processes and mechanisms underlying successful biological invasions.

## Introduction

Invasive species are a global problem, the economic costs of which total in the tens to hundreds of billions of (US) dollars each year (Pyšek & Richardson, 2010; Early *et al*., 2016). Yet the basic question has yet to be answered: Why do some introduced species become invasive, and others not? Attempts to answer this pivotal question have primarily focused on ecological explanations rather than the evolutionary processes acting during invasions. Notably, over 30 “invasion hypotheses” have been formulated to explain invasion success, many of which are not mutually exclusive (Enders *et al*., 2020). These include: propagule pressure (i.e., number and/or volume of introductions; Lockwood *et al*., 2005; Vahsen *et al*., 2018); the evolution of increased competitive ability and related enemy release hypothesis (Blossey & Nötzold, 1995); habitat filtering (Weiher & Keddy, 1995); rapid post-introduction evolution (Prentis *et al*., 2008; Colautti & Barrett, 2013; Felker-Quinn *et al*., 2013); and the “ideal weed” hypothesis (Baker, 1965; Rejmánek & Richardson, 1996).

In the last two decades, we have witnessed an explosion in the application of genetic and genomic tools in invasion biology, leading to reassessment of the evolutionary processes underlying invasions. Recent examples in invasive plants include genomic studies of the globally invasive common ragweed (van Boheemen *et al*., 2017; Battlay *et al*., 2023), yellow starthistle (Turner *et al*., 2013; Barker *et al*., 2017), white clover (Innes *et al*., 2022; Battlay *et al*., 2025; Hendrickson *et al*., 2026), and weedy rice (Qiu *et al*., 2020; Presotto *et al*., 2024). The picture emerging from these studies shows that invasions are often more complex than previously thought, including multiple introductions, long-distance dispersal, admixture between historically isolated populations, and rapid post-introduction evolution. Thus, examples supporting the “traditional” hypothesis of invasions as singular events, resulting in extreme genetic bottlenecks (i.e., the “genetic paradox” of invasions) seem to be in the minority (but see Pascual-Díaz *et al*., 2025; Rizzieri *et al*., 2026).

The genetic composition of introduced populations is shaped by complex interactions among colonization history, genetic drift, expansion, admixture, and selection (Dlugosch & Parker, 2008; Keller & Taylor, 2008; Dlugosch *et al*., 2015; Sherpa & Després, 2021; Hodgins *et al*., 2025). While genetic bottlenecks are expected to constrain evolutionary potential in introduced populations (Nei *et al*., 1975), multiple introductions followed by admixture can facilitate establishment and spread by alleviating such bottlenecks (Dlugosch & Parker, 2008; Barker *et al*., 2019), increasing the fitness of hybrids due to heterosis (Keller *et al*., 2014; Vercellino *et al*., 2023), and creating novel genetic combinations on which selection can act (Ellstrand & Schierenbeck, 2006; Presotto *et al*., 2024; Hernández *et al*., 2024). For these reasons, admixture between multiple introductions has been proposed as a general mechanism to explain invasion success despite genetic bottlenecks (Verhoeven *et al*., 2011; Bock *et al*., 2015; Dlugosch *et al*., 2015). On the other hand, multiple introductions themselves can facilitate invasions by increasing the chances that pre-adapted genotypes find suitable climates in the introduced range (i.e., habitat filtering; Gamba *et al*., 2025), and admixture can also prevent adaptation by breaking up favorable allele combinations (Verhoeven *et al*., 2011; Barker *et al*., 2019). Resolving the timing and relative contribution of distinct evolutionary processes is thus critical to understanding the role each process plays in invasion success.

Herbarium genomics, here defined as the study of genome-wide sequences from preserved historical plant specimens, holds great potential for reconstructing invasion histories by allowing the direct observation of genotypes from distinct stages of the invasion timeline (Lang *et al*., 2019; Kreiner *et al*., 2022; Bieker *et al*., 2022; Battlay *et al*., 2023, 2025; Kim *et al*., 2023; Hendrickson *et al*., 2026). Historical specimens can reveal cryptic colonization events that have been replaced in the present (Rawlence *et al*., 2017; Hendrickson *et al*., 2026), and can provide direct information on the timing, magnitude, and persistence of genetic bottlenecks (Chan *et al*., 2006; Wales *et al*., 2019) as well as on the timing and magnitude of admixture (Muhlfeld *et al*., 2014; Kreiner *et al*., 2022). Thus, studying historical specimens enables us to connect genetic patterns to processes in invasion biology.

A powerful approach for cost-effectively interrogating whole genomes of historical collections is through low-coverage whole-genome sequencing (lcWGS; reviewed in Lou *et al*., 2021; Lou & Therkildsen, 2022). This approach requires only basic DNA library preparation and can accommodate both contemporary and historical samples (Hendrickson *et al*., 2026). Indeed, sampling more individuals at low coverage depth can yield estimates of many key population metrics at equal or greater accuracy than sampling fewer individuals at higher depth (Alex Buerkle & Gompert, 2013; Fumagalli, 2013; Lou *et al*., 2021; Lou & Therkildsen, 2022). Therefore, lcWGS can effectively overcome both the low resolution of reduced-representation sequencing (Lowry *et al*., 2017) and the prohibitive cost of high-coverage whole-genome sequencing.

Well-established invasions, while having low probabilities of eradication, can be leveraged as powerful and informative systems to study the spatiotemporal dynamics of the biological invasion process, ultimately leading to more refined detection and control efforts that can be utilized for new invaders (North *et al*., 2021). Here we focus on stiltgrass (*Microstegium vimineum* [Trin.] A. Camus), an invasive C_4_ grass species introduced to eastern North America over a century ago (Fairbrothers & Gray, 1972), and now established in 31 USA states and Ontario, Canada (EDDMapS; https://www.eddmaps.org, accessed 18 November 2025). Stiltgrass has a mixed mating system, with terminal wind-pollinated chasmogamous flowers that permit outcrossing and axillary cleistogamous flowers that self-pollinate. Individual plants can produce over 1,000 seeds per year, and seed can persist in the seed bank for up to five years and even longer under dry conditions (Cheplick, 2010; Redwood *et al*., 2018). It is a putative paleopolyploid with 2*n* = 20. Its chromosome-scale genome shows a 2:1 syntenic relationship with *Sorghum* and 30% duplicated complete BUSCOs, consistent with a lineage-specific whole-genome duplication followed by extensive diploidization and chromosomal rearrangement. Available *K*_s_ evidence places this duplication within approximately the last 10 Myr, whereas the inferred 1–2-Myr event represents an expansion of LTR retrotransposons (Ramachandran et al., 2021).

Stiltgrass is rapidly becoming a model species for understanding the evolutionary genomics of invasions. The first specimen in North America was collected in 1919 near Knoxville, Tennessee, USA (Fairbrothers & Gray, 1972). Stiltgrass was thought to be introduced to North America as porcelain packing material from the native range in eastern Asia (Tu, 2000). Regular imports of porcelain presumably resulted in numerous introductions of seeds attached to the stems used as packing material, thus creating multiple human-assisted opportunities for invasion. By analyzing over 1,000 digitized herbarium records in the native and introduced ranges, and based on the presence/absence of awns, Barrett *et al*. (2022) hypothesized that at least two separate invasions occurred in North America: an awnless form in the southern USA and the awned form in the northern USA. A short-awned form that is found in geographically intermediate populations may have arisen via secondary contact after spread from introduction points. This hypothesis includes three predictions: 1) there are at least two introductions to North America; 2) there is substantial admixture that leads to the formation of novel phenotypic variation; and 3) selection has rapidly occurred on awns during invasion.

Here we investigate how colonization, admixture, and selection shaped the North American invasion of stiltgrass. We leverage low-coverage whole-genome sequencing of historical and contemporary samples spanning the temporal and spatial extent of the North American invasion and the native range in East Asia to address four questions. First, are there multiple colonization events and if so, do they come from different source populations? Second, are colonization events accompanied by strong genetic bottlenecks? Third, given the presence of multiple colonization events, is there evidence of admixture following secondary contact that increases genetic diversity? Fourth, do temporal genomic scans reveal signatures of post-introduction selection? Our results provide evidence for a complex invasion scenario that includes multiple introductions, limited admixture, and selection acting on a rapid timescale.

## Materials and methods

### Sampling

Field-collected samples were acquired between the years 2019–2022, hereafter referred to as contemporary, whereas herbarium samples, originally collected between 1885 and 2013, are referred to as historical. For contemporary sampling, we collected 209 individuals from 31 localities in the USA, 56 from six localities in China, 49 from nine localities in Japan, and 16 from three localities in Taiwan (Table **S1**; Fig. **1a**). Three leaf blades were sampled from each plant collected in the field, placed in a coin envelope, then preserved with silica gel desiccant in an airtight container and stored at room temperature until DNA extraction. A single plant from each locality was pressed and dried as a voucher specimen and deposited at WVA. For historical samples, 113 were from the USA, and 29 were from Asia. Historical sampling was conducted at the following US herbaria: BH, BRIT, CM, MO, OS, PH, TENN, US, and WVA (Table **S1**). These included the first US historical collection of stiltgrass from Knox County, Tennessee (1919). Historical collections from Asia ranged from 1885–1998 (native range) and 1919–2013 for the USA (introduced range). For historical samples, approximately 2 cm^2^ of leaf tissue was collected either from the attached packet or the specimen sheet, the latter only if the amount of material on the sheet was deemed sufficient by herbarium staff as not to reduce the utility of the specimen. All specimens were annotated according to institutional policies. Herbarium-derived leaf tissues were similarly placed in coin envelopes, then stored in silica gel desiccant before DNA extraction. Taxonomic screening led to the exclusion of 32 contemporary samples collected in China and nine historical samples collected in China, Japan, India, and Taiwan (Table **S1;** Supplementary information) from downstream population genomic analyses, leaving 431 samples: 298 contemporary and 133 historical.

**Figure 1.**
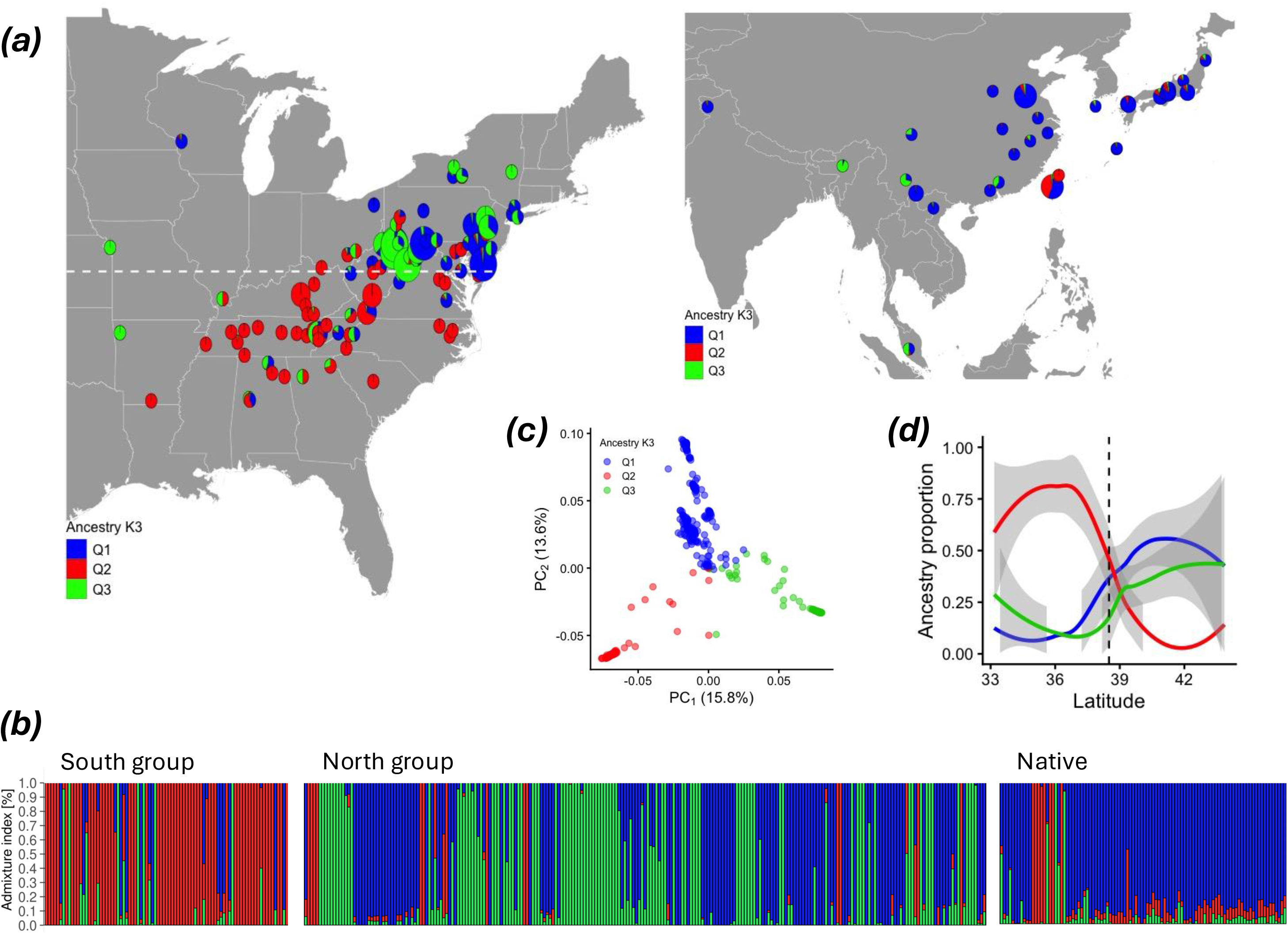
Population structure in the native and introduced ranges. (a) The geographic distribution of clusters inferred from *NGSadmix* in the introduced range (left) and the native range (right). (b) Ancestry proportions at K = 3. Samples are sorted by latitude from lowest to highest. (c) Principal component analysis colored by the largest ancestry group from *NGSadmix*. (d) Latitudinal cline of ancestry proportions in the introduced range.

### DNA extraction and sequencing

Total genomic DNA was extracted with the CTAB method (Doyle & Doyle, 1987), using approximately 1-2 cm^2^ of leaf tissue, with scissors sterilized with 100% ethanol for both field-and herbarium-derived tissues. DNA pellets were eluted in 70-120 µl Tris-EDTA buffer (TE, pH 8.0). For herbarium extractions, an additional chloroform-isoamyl alcohol (24:1 volume:volume ratio) extraction step was conducted. DNA samples were run individually on 1% agarose gels to visualize DNA quality and the degree of post-mortem shearing, and DNA concentration was quantified with the Qubit Broad Range dsDNA assay (Thermo Fisher Scientific, Waltham, Massachusetts, USA). All extractions were conducted on a lab countertop sterilized with sodium hypochlorite (10% bleach solution) followed by sterilization with 100% ethanol. DNA samples were diluted to 20 ng µl^-1^ with TE and stored at -80 °C. Illumina sequencing libraries were prepared with the SparQ DNA Frag and Library kit (Quantabio, Beverly, Massachusetts, USA), using two-fifths of the recommended reaction volume. The suggested 14-minute fragmentase time (used for all contemporary samples) was reduced to a one-minute time for herbarium-derived DNA that showed evidence of degradation (i.e., the predominance of low-molecular-weight DNA as assessed by gel electrophoresis), to prevent over-shearing while still preparing the pre-fragmented libraries for adapter ligation. Otherwise, library preparation followed the manufacturer’s protocol. Libraries were quantified with the Qubit High Sensitivity dsDNA assay. Libraries of low concentration (i.e., less than 2 ng ul^-1^) were amplified with the Kapa Library Amplification kit (Roche, Basel, Switzerland) with 12-18 PCR cycles following the kit instructions, and re-quantified.

Libraries were pooled at equimolar ratios and fragment sizes were visualized with an Agilent Bioanalyzer (Agilent, Santa Clara, California, USA) before final pool quantification via quantitative PCR at the WVU Genomics Core Facility. If needed, pools were run on a Blue Pippin Prep (Sage Science, Beverly, Massachusetts, USA) to remove any adapter dimer carried over from library preparation and re-quantified as above. Sequencing was conducted for pools of 75-90 accessions on six lanes of an Illumina NextSeq2000 sequencer at the WVU-Marshall University Genomic Core Facility using 200-cycle P3 chemistry, resulting in approximately 10^9^ 2 × 100-bp paired-end reads per run and targeting approximately 1.0–1.2 × 10^7^ read pairs per accession.

### Data processing and read mapping

Reads were processed with *fastp v.0.23.4* (Chen et al., 2018) with the following options (--cut_right --dedup -g) to quality-trim and trim polyG tails using a sliding window approach. Pre- and post-processed read qualities were assessed with *FastQC* (www.bioinformatics.babraham.ac.uk/projects/fastqc/). Processed reads were mapped to the reference genome (Ramachandran *et al*., 2021) using *BWA-MEM v.0.7.18* (Li & Durbin, 2009). Mapping statistics for each sample, including mean coverage, were estimated using *BamTools v.2.5.2* (Barnett *et al*., 2011) and *Qualimap v.2.3* (Okonechnikov *et al*., 2016). We also used *mapDamage v.2.0* (Jónsson *et al*., 2013) to quantify DNA damage patterns on historical samples.

Mapped read coverage depth ranged from 0.0002 to 10.5x (mean = 1.66x) and was similar between historical and contemporary samples (mean = 1.76x and 1.62x, respectively; t-test *P* = 0.4088). To reduce differences in coverage among samples, for downstream analyses we down-sampled coverage to 2.5x for any individual with > 2.5x coverage (N = 45).

Historical samples showed characteristic ancient-DNA damage: C-to-T misincorporation at the first base ranged from 0.16% to 2.37% (mean = 1.01%), significantly more than in contemporary samples (range = 0.05-1.41%; mean = 0.26%; t-test *P* < 2.2 × 10^-16^), but similar between ranges (mean = 1.13% and 0.98% for native and introduced ranges, respectively; t-test *P* = 0.1706). The C-to-T base misincorporation at the first base decreased with the year of collection in both native and introduced ranges (slope = -6.44 × 10^-5^ and -3.49 × 10^-5^, respectively; *P* < 0.001 for both). Therefore, to account for potential artifacts from ancient-DNA damage, in comparisons between historical and contemporary samples, we also estimated genetic diversity after removing transitions (i.e., A↔G and C↔T; the -rmTrans 1 *ANGSD* option).

### Genotype likelihood estimation and SNP filtering

Genotype likelihoods for each site and individual were estimated using *ANGSD v.0.941* (Korneliussen *et al*., 2014) with the following options [-nInd n -minInd 0.3*n -doMajorMinor 1 -doPost 1 -doGlf 2 -doMaf 1 -gl 2 -minMapQ 30 -minQ 20 -minMaf 0.05 -SNP_pval 1e-6 -r CHR -baq 1 -uniqueOnly 1 -only_proper_pairs 1 -ref REF], with n being the number of samples, REF the reference genome in fasta format, and CHR each of the 23 largest scaffolds. We then estimated pairwise linkage disequilibrium (LD) among SNPs within a 50-kb distance using *ngsLD* (Fox et al., 2019) and pruned SNPs with r^2^ > 0.2. The code for all analyses is available at: https://github.com/fernandoh33/Invasion-genomics-stiltgrass.

To calculate genetic diversity and neutrality statistics, we estimated the site allele frequency likelihoods (.saf file) using all variant and invariant sites with the following options [-nInd n -minInd 0.3*n -doMajorMinor 4 -doPost 1 -doGlf 2 -doMaf 4 -baq 1 -minMapQ 30 -minQ 20 -anc REF -ref REF -r CHR] for each group. For estimating global and window-based pairwise F_ST_ between groups, we estimated 2D folded site frequency spectrum files using all variant sites, adding the options: -SNP_pval 1e-6 and -minMaf 0.01.

### Genetic diversity and population structure

Principal component and admixture analyses were performed with *pcangsd v.1.36.4* (Meisner & Albrechtsen, 2018) and *ngsadmix v.0.939* (Skotte et al., 2013), respectively, using the filtered Beagle files with genotype likelihoods generated above with no additional filters. For *ngsadmix*, we ran 10 independent replicates using distinct random seeds for K = 1 – 10. The optimal K was determined using the Evanno method (Evanno et al., 2005) implemented i*n CLUMPAK v.1.1* (Kopelman et al., 2015). For each K, we used the most likely run for estimating ancestry proportions for each sample and for visualization. The admixture index for each sample was estimated as 1 minus the maximum ancestry proportion of that sample.

Genetic diversity and neutrality statistics (Watterson’s θ, π, and Tajima’s D) and pairwise F_ST_ between groups were estimated for non-overlapping 10-kb windows as well as for each of the 23 largest scaffolds. The median values of scaffold-based estimates hereafter are referred to as global. For window-based estimates of genetic diversity and neutrality statistics, only windows with at least 1,000 sites and present in at least 30% of individuals were included. For pairwise F_ST_, only 10-kb windows with at least 5 SNPs were included.

### Geographic source of invasive populations

We used *locator v.1.1* (Battey *et al*., 2020) to identify the putative geographic sources of introduced populations. As *locator* cannot handle genotype likelihoods, we called SNPs with *bcftools* v.1.19 (Danecek *et al*., 2021) in a subset of samples with coverage > 2x (n = 149, n_NATIVE_= 45, n_INVASIVE_= 104). For analyses, we randomly selected 10,000 SNPs from a set of hard-filtered SNPs with mean base quality > 30, mean mapping quality > 20, mean depth between 1 and 10, minimum allele frequency > 0.1, low linkage disequilibrium (r^2^ < 0.2), and missingness < 0.5. Native and introduced samples were used as training and test populations, respectively. We generated 100 random seed numbers and ran *locator* 100 times. Instead of using a single best model, we retained all models with prediction accuracies (r^2^ between actual and predicted values in the native range) > 0.99 for either latitude or longitude. Then, for each sample, we estimated latitude as the median of all models with r^2^ > 0.99 and repeated this procedure for longitude. We repeated this analysis with two additional datasets, a low differentiation dataset with SNPs within regions of low differentiation between awned and awnless phenotypes in the introduced range (F_ST_ < 0.1), and a “divergent” dataset with SNPs from the top 1% windows of differentiation (F_ST_ > 0.26).

### Demographic modelling

We performed demographic modelling with *fastsimcoal v.2709* (Excoffier et al., 2021) with two objectives: 1) evaluating whether divergence between distinct southern and northern groups (see results) occurred before or after introduction to the USA; and 2) estimating demographic parameters (population size, bottleneck intensity, post-introduction expansion, and timing of introduction) that characterize the invasion process. For demographic models, we filtered sites to retain variant and invariant four-fold degenerate sites, which were identified using *degenotate v.1.3* (Mirchandani *et al*., 2024), produced multidimensional derived allele site-frequency spectrum files with *ANGSD* (Korneliussen *et al*., 2014) using an ancestral reference genome, and assumed a mutation rate of 1 × 10^-9^ per site per generation and a generation time of 1 year.

For generating the ancestral reference genome, we mapped reads of 29 samples of *M. fasciculatum* (Supplementary information) from Yunnan and Guangdong onto the *M. vimineum* reference with the following options [-doFasta 2 -minMapQ 30 -minQ 20 -baq 1 -ref REF - doCounts 1 -domaf 1 -domajorminor 1 -gl 2 -minind 10 -explode 1]. Because this reference includes variant sites, we identified SNPs in these samples adding the option -SNP_pval 1e-6 in a separate analysis and then masked the reference using the maskfasta option in *bedtools* v.2.31.1 (Quinlan & Hall, 2010). The resulting ancestral reference includes 243,178,637 invariant sites and 4,559,081 4-fold degenerate sites (approximately 20% of all reference sites and 50 % of four-fold degenerate reference genome sites, respectively).

For models estimating divergence time among genetic groups, we assumed no gene flow and constant population sizes along each branch. We estimated seven parameters: current population size for each genetic group (*N_Q1_*, *N_Q2_*, and *N_Q3_* for North Q1, South Q2, and North Q3, respectively), divergence time between North Q1 and South Q2 (*T_Q1-Q2_*) and between the ancestral population Q1-Q2 and North Q3 (*T_Q12-Q3_*), and ancestral population size for the ancestral populations Q1-Q2 and Q1-Q2-Q3 (*N_Q12_* and *N_Q123_*, respectively). We tested three alternative tree topologies. Each model was run 100 times with 100 cycles (-L 100) and 100,000 simulations (-n 100,000), and runs were compared using the Akaike Information Criterion (AIC). To directly test whether genetic groups in the introduced range underwent genetic bottlenecks during introductions to the USA, we ran models for each group separately and estimated six parameters: current population size (*N_CON_*), timing of bottleneck (*T_SBOT_* and *T_EBOT_* for the start and end of the bottleneck, respectively), growth rate after the end of the bottleneck, population size during the bottleneck (*N_BOT_*), and ancestral population size (*N_ANC_*). Bottleneck duration was estimated as (*T_SBOT_* - *T_EBOT_*) and bottleneck intensity as (*N_BOT_ / N_ANC_*). Times are measured in years before present, so T_SBOT_ is older than T_EBOT_.

All scripts and input files used for demographic modelling can be found at: https://github.com/fernandoh33/Invasion-genomics-stiltgrass.

### Temporal patterns of population structure and genetic diversity

We performed a time series analysis to investigate temporal changes in population structure and genetic diversity. We sorted historical samples by collection year and used a sliding-window approach (window = 10 samples, steps = 5 samples) to track changes in Watterson’s θ, nucleotide diversity (π), and Tajima’s D over time. We hypothesized that multiple introductions alleviate genetic bottlenecks, resulting in significant relationships between time and θ and π (positive relationship) and/or Tajima’s D (negative relationship). The sliding window approach resulted in Watterson’s θ, π, and Tajima’s D estimates for 10 temporal windows for each South (samples from latitude < 38.5°) and North (samples from latitude > 38.5°) groups and 22 temporal windows when all historical samples from the introduced range were combined in a single group. Genetic statistics were estimated for each scaffold and temporal window. For each window, we estimated the median year of collection and fitted a loess model for θ, π, and Tajima’s D with a smoothing span of 0.7 using the geom_smooth function in *ggplot2 v.4.0.3* (Wickham, 2016).

### Genomic signatures of selection following colonization

We investigated temporal signatures of selection by comparing historical and contemporary samples. Based on the temporal distribution of sampling and the temporal patterns of genetic diversity (see results), we divided samples collected in the introduced range into three temporal slices: historical samples collected before 1960 (hereafter before 1960), historical samples collected between 1960 and 2013 (hereafter after 1960), and contemporary samples, collected in the field between 2019 and 2022. All samples with the awnless phenotype and latitude < 38.5° were assigned to the South group, while all samples with the awned phenotype and latitude > 38.5° were assigned to the North group. We used 38.5° as the inflection point because it marks the division between the USA awned and awnless forms in digitized herbarium samples in previous research (Barrett *et al*., 2022) as well as in our own study. To avoid biases in allele frequency estimations, for two populations with more than five individuals (collected in Alleghany County, Virginia, USA and Monongalia County, West Virginia, USA, both in the North group), we randomly selected five individuals with at least 1x coverage of mapped reads. After applying the above criteria, we retained 24, 28, and 35 samples for the South groups before 1960, after 1960, and contemporary, respectively, and 25, 25, and 34 samples for the North groups before 1960, after 1960, and contemporary, respectively. We also compared introduced groups with native populations, clustering native samples by country: China, Japan, and Taiwan. For China, due to the low number of contemporary samples after excluding samples from Guangdong and Yunnan (see phylogenetic analysis; Fig. S**1**), historical samples were included to increase sample size to n = 22. For Japan, we used only contemporary samples and retained up to five individuals per site (n = 37), while for Taiwan, all 16 contemporary samples were included.

We identified putative selective sweeps in the North and South separately as windows showing both extreme genetic differentiation and an extreme reduction in genetic diversity over time, i.e., windows above the empirical 95^th^ percentile of both temporal F_ST_ and ln(π_B_/π_C_) distributions, with π_B_ being the estimate for samples collected before 1960 and π_C_ the estimate for contemporary samples. Only windows with at least five SNPs for F_ST_ comparisons and 1,000 sites in both groups under comparison for π estimates were included in the analysis. After applying these criteria, we retained 100,920 of 110,842 and 96,188 of 110,864 windows for the North and South comparisons, respectively. For candidate outlier regions, we extracted gene annotations from the outlier window and the immediately adjacent windows.

## Results

### Genetic diversity and population structure

There were distinct patterns of population structure across both the native and introduced ranges of *M. vimineum* (Fig. **1a**). *NGSadmix* supported the presence of three main genetic clusters (Fig. **1b**), and these clusters were largely separated in the principal component analysis (Fig. **1c**). In the native range, most samples were assigned to the Q1 group (96 out of 109), though several contemporary samples from Taiwan were assigned to the Q2 group and four samples (three historical samples from China, Malaysia and India, and one contemporary sample from Taiwan) were assigned to Q3 (Fig. **1a**; Table **S1**). In the introduced range, genetic groups corresponded closely with geography and with awn phenotypes. Samples from the southern region (latitude < 38.5°) presented the awnless phenotype (80/82 samples) and were assigned to the Q2 genetic group (63/85 samples; Fig. **1a,b,d**). Samples from the northern region (latitude > 38.5°) presented the awned phenotype (123/175 samples) and were assigned to either the Q1 or Q3 groups (220/237 samples; Fig. **1a,b,d**).

Population structure was lower in the native (F_ST_ = 0.07, 0.06, and 0.12 for China-Japan, China-Taiwan, and Japan-Taiwan, respectively) than in the introduced range (F_ST_ = 0.18, 0.43, and 0.28 for South Q2 - North Q1, South Q2 - North Q3, and North Q1 - North Q3, respectively). In addition, North and South groups in the USA were genetically less differentiated from native populations than from each other (mean F_ST_ = 0.16 and 0.29, respectively).

### Multiple independent introductions underlie colonization of North America

The trained networks implemented in *locator* support at least two independent introductions in the introduced range: Japan for North Q1 and South Q2, and China for North Q3 (Fig. **2a,b**). No samples were predicted to be introduced from Taiwan, although we only have three samples from Taiwan in the training dataset. Six samples (out of 104) showed intermediate locations, possibly reflecting post-introduction admixture or assignment uncertainty.

**Figure 2.**
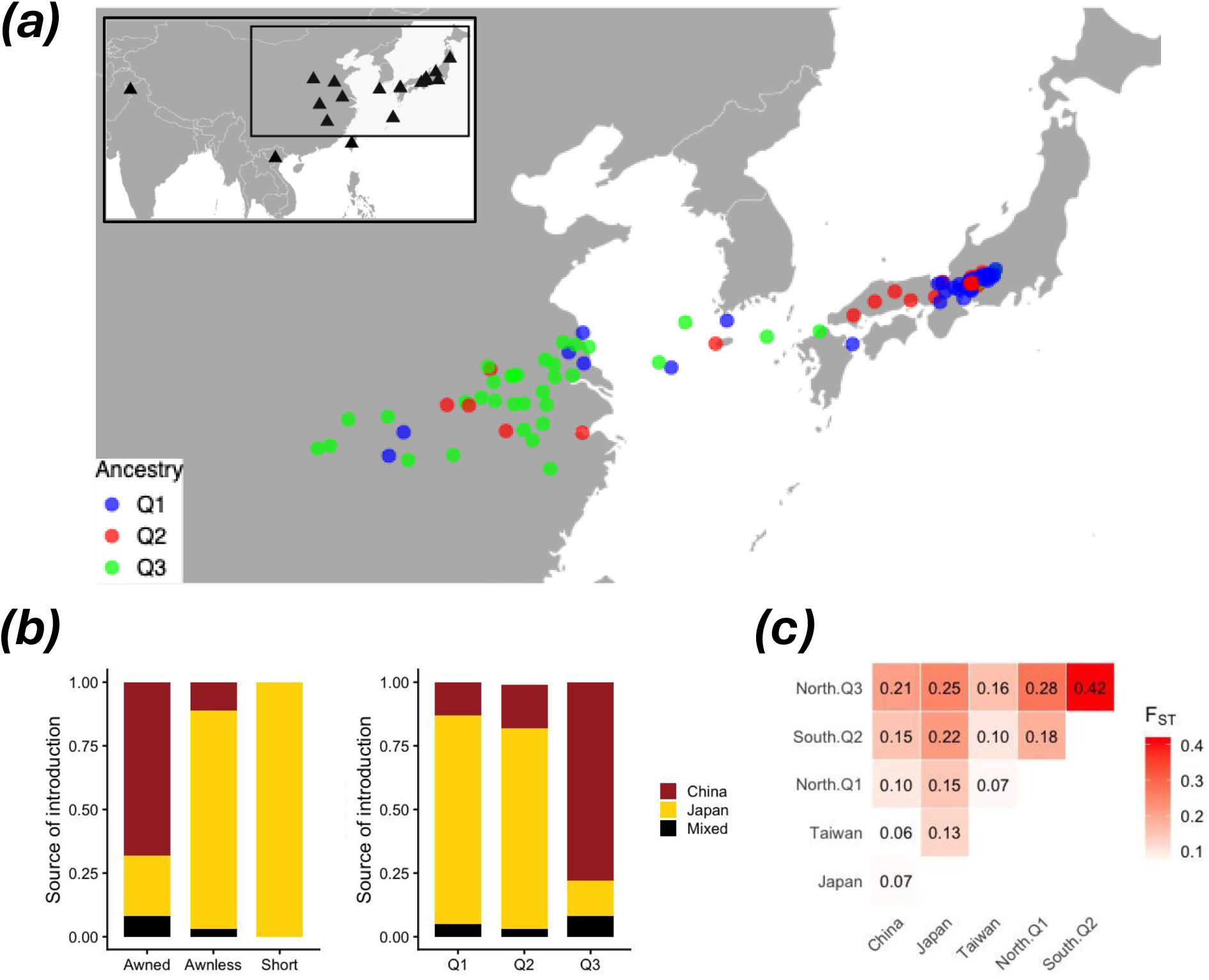
Geographic source prediction. (a) Predicted location of introduced samples in the native range. Dots in the main plot represent samples from the introduced range colored by genetic groups inferred from *NGSadmix* (Fig. 1). Black triangles in the inner plot represent native locations used in the training dataset. (b) Bar plots showing predicted introduction source by awn phenotype (left) and genetic group (right). (c) Pairwise F_ST_ between native and introduced groups. Each value is the median of estimates for the 23 largest scaffolds.

The awn phenotype was also strongly associated with the introduction source; most samples with the awnless phenotype were predicted to be introduced from Japan (31 out of 36), while most samples with the awned phenotype were predicted to be introduced from China (26 out of 38) (Fig. **2b**). For all samples with the short awn phenotype (n = 12), Japan was the putative source of introduction. To evaluate whether genomic regions associated with the awn phenotype were biasing the identification of the geographic source, we ran locator with two additional datasets: a low-differentiation dataset with SNPs within regions of low differentiation between awned and awnless phenotypes in the introduced range (F_ST_ < 0.1), and a divergent dataset with SNPs within the top 1% windows of differentiation (F_ST_ > 0.26). We found similar patterns for all three datasets with high correlation between low-differentiation and divergent models (r^2^_LAT_ = 0.91; r^2^_LON_ = 0.88).

We also evaluated whether the inferred pairwise F_ST_ values supported the geographic source of introduction. Source prediction was not fully supported by pairwise F_ST_ comparisons (Fig. **2c**); the three introduced groups were most closely related to Taiwan (Fig. **2c**), followed by China and then Japan. Together, *locator* and pairwise F_ST_ results suggest that multiple introductions acted as a major driver of phenotypic and geographic differentiation in the introduced range, though the geographic source of independent introductions is unclear.

### Temporal patterns of population structure and genetic diversity

For contemporary samples, genetic diversity was higher in the native range than the introduced range (median Watterson’s θ = 0.0084 and 0.0074; median π = 0.0051 and 0.0041, respectively), suggesting an overall weak genetic bottleneck in the introduced range. Within the introduced range, however, the northern region (North Q1 and North Q3 groups combined) showed much higher genetic diversity than the southern region (Watterson’s θ = 0.007 and 0.0031; π = 0.0042 and 0.0026, respectively). In addition, the North Q1 group showed higher genetic diversity than the North Q3 group (Watterson’s θ = 0.0056 and 0.0025; π = 0.0041 and 0.0014, respectively).

Time series analysis showed that all three genetic groups were present in the introduced range before 1950 (Fig. **3a**), and admixture between clusters was relatively low in historical as well as contemporary samples (Fig. **3b**). Genetic diversity was highest before 1960, intermediate after 1960, and lowest in contemporary samples in both northern and southern regions (Fig. **3c**). The sliding-window analysis also showed a temporal decline in diversity (Fig. 3c). The reduction in nucleotide diversity between pre-1960 and contemporary samples persisted after transitions (A↔G and C↔T) were removed (Fig. S2). Tajima’s D was lowest before 1960 in both regions (Fig. **3c**), consistent with rapid range expansion following a bottleneck. After 1960, Tajima’s D remained low in the southern region but increased in the northern one. Contemporary values were generally less negative than those in the earliest historical group, although median values remained below zero in both regions.

**Figure 3.**
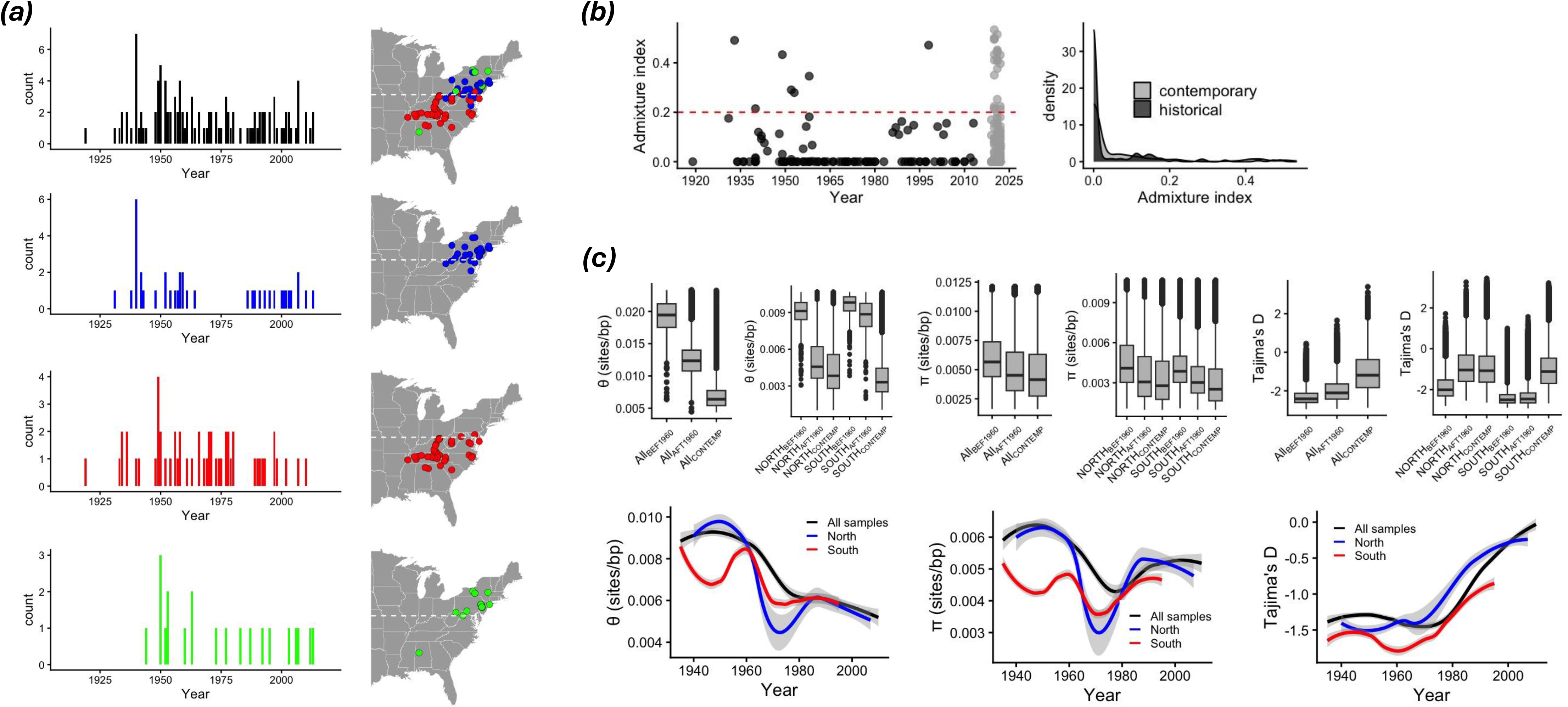
Temporal patterns of population structure and genetic diversity in the introduced range. (a) Temporal and geographic distribution of all historical samples (upper plot) and individual genetic groups (bottom plots). Genetic groups were inferred from *NGSadmix* (Fig. 1). (b) Temporal patterns of admixture index over time in historical (black dots) and contemporary samples (gray dots). The admixture index for each sample was estimated as 1 minus the maximum ancestry proportion of that sample. (c) Genetic diversity and neutrality statistics over time. Upper plots: boxplots compare samples collected before 1960, after 1960 (between 1960–2013) and between 2019–2022 (contemporary), with the introduced range pooled and with the North and South shown separately. Lower plots: sliding-window analysis of historical samples (window = 10 samples, step = 5 samples) for Watterson’s θ, nucleotide diversity (π) and Tajima’s D. For each window, we estimated the median year and fitted a loess model for all samples (black line) as well as north (blue line) and south (red line) groups separately with a smoothing span of 0.7.

Demographic modelling indicates that divergence among genetic groups occurred well before the introduction to the USA (Fig. **4a**). In the best model (Model 3), all three groups diverged from a common ancestor 58,563 y ago (Table **S2**). Similar divergence times were found for the best run of Model 1 (*T_Q1-Q2_* = 33,570 y; *T_Q12-Q3_* = 44,540 y; Table **S2**) and Model 2 (*T_Q1-Q3_* = 27,067 y; *T_Q13-Q2_* = 57,324 y; Table **S2**), supporting deep divergence among groups across the demographic models considered and suggesting that divergence did not occur following introduction.

**Figure 4.**
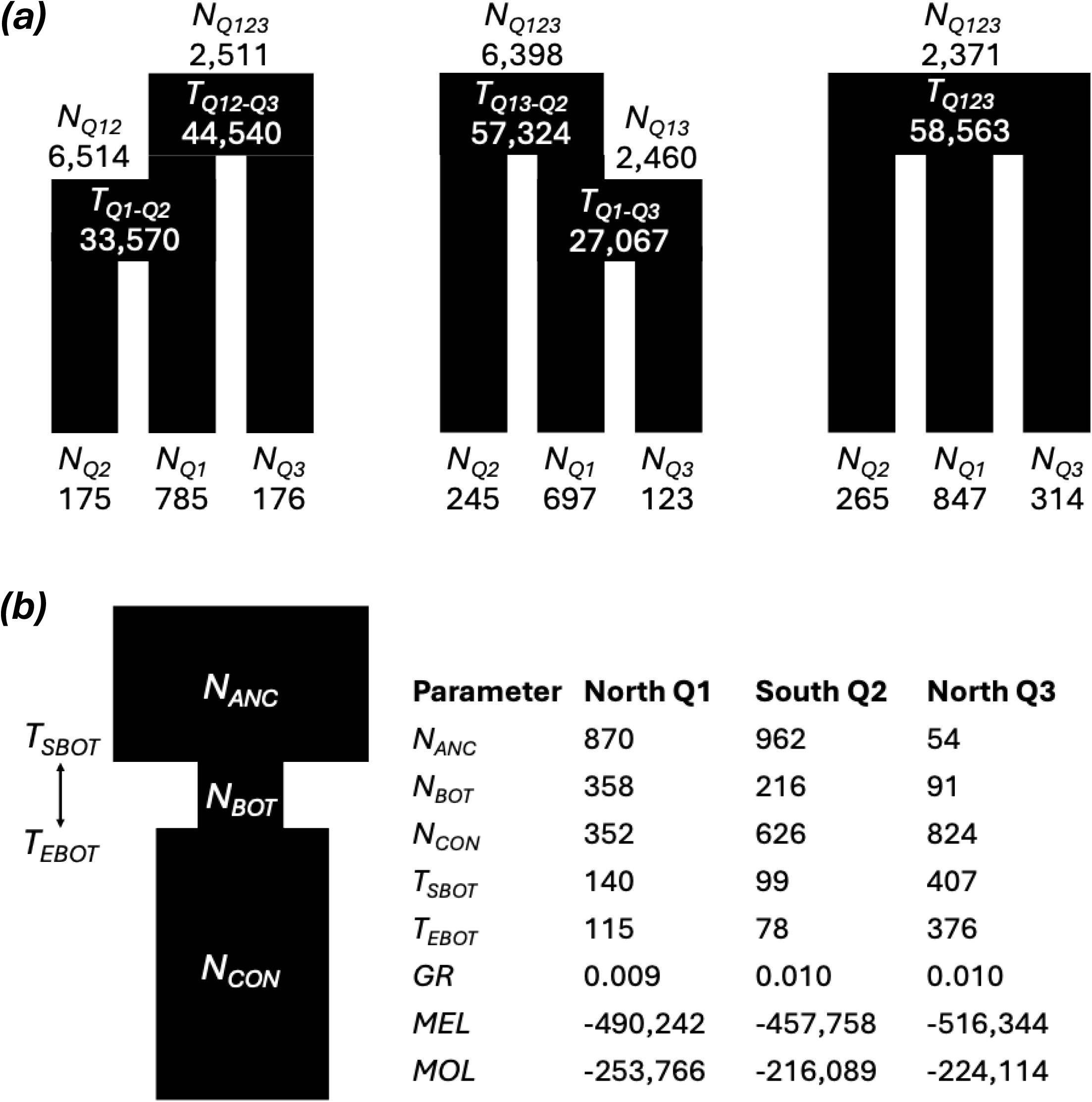
Demographic modelling in *fastsimcoal2*. (a) Schematic diagrams of the three divergence models compared to estimate population size and divergence time of genetic groups and their ancestral populations. *N_Q1_, N_Q2_,* and *N_Q3_* are extant population sizes of genetic groups Q1, Q2, and Q3, respectively while *N_Q12_, N_Q13_*, and *N_Q123_* are population sizes of the ancestral population of Q1 and Q2, Q1 and Q3, and all genetic groups, respectively. *T_Q1-Q2_*, *T_Q1-Q3_, T_Q12-Q3_*, *T_Q13-Q2_* and *T_Q123_* are divergence times between Q1 and Q2, Q1 and Q3, the ancestor of Q1-Q2 and Q3, the ancestor of Q1 - Q3 and Q2, and all populations, respectively. For each model, parameters for the best run are provided. (b) Schematic diagram of the model run separately for each group, and parameters for the best run of each model. *N_ANC_*, *N_BOT_*, and *N_CON_* are population sizes before and during the bottleneck, and at the present, respectively. *T_SBOT_* and *T_EBOT_* are the times in years before present at the start and end of the bottleneck and GR is growth rate after the bottleneck. *MEL* and *MOL* are maximum estimated and observed base-10 log-likelihoods. Population sizes are shown in thousands of haploid individuals; unscaled estimates are provided in Tables S2 and S3. Times are in years before present.

The best-fitting demographic models inferred substantial, short-lived population contractions near the period of introduction to the USA in two of the three genetic groups. For North Q1 and South Q2, the best runs indicate substantial population contractions (*N_BOT_/N_ANC_* = 0.41 and 0.22) beginning 140 and 99 y ago, and lasting 25 and 21 y, respectively (Fig. **4b**; Table **S3**). For North Q3, the best run indicates an increase in population size during the interval 407–376 y ago (Fig. **4b**; Table **S3**), whereas five of the ten highest-likelihood runs inferred more recent contractions (*T_SBOT_* = 78 – 104 y; *T_EBOT_* = 64 – 98 y; *N_BOT_* = 5,013 – 735,241; Table **S3**). These five runs had lower likelihoods than the highest-likelihood solution.

### Temporal patterns of selection

We compared genome-wide genetic differentiation between historical and contemporary samples within the northern (Q1 and Q3 combined) and southern regions. Genome-wide, temporal F_ST_ showed little to no differentiation in the northern or southern regions (all temporal genome-wide F_ST_ < 0.02), indicating lineage persistence after initial introductions (i.e., genetic continuity). By contrast, genetic divergence between southern and northern regions increased over time (F_ST_ = 0.16, 0.23, and 0.35 for comparisons before 1960, after 1960, and for contemporary samples, respectively), suggesting that natural selection and/or genetic drift post-introduction may have contributed to observed patterns of differentiation in the introduced range.

We then scanned the genome to identify putative selective sweeps, observed as outlier windows showing both extreme genetic differentiation between historical and contemporary samples as well as an extreme reduction in genetic diversity over time (Fig. **5a**). We identified 226 and 478 outlier 10-kb windows in the northern and southern regions, respectively; these windows and their immediate neighbours contained 329 and 609 genes (Fig. **5b**). When we compared the outlier genes from each region, we found relatively few shared genes but more than expected by chance (12 overlapping genes; hypergeometric test, *P* = 2.1 × 10^−3^, expected number = 4.5). Among these overlapping genes, we found two putatively associated with flowering time: EARLY HEADING DATE 2 (*Ehd2*) on scaffold 1, and PHOTOPERIOD-INDEPENDENT EARLY FLOWERING 1 (*PIE1*) on scaffold 4. Within each region, we also identified genes putatively associated with: (1) flowering time: EARLY FLOWERING 4 (*ELF4*) and CONSTANS-like (*CO*) in the North; (2) seed dormancy: SUPPRESSOR OF ABI3-5 (*SUA*) in the South; and 3) plant development: WUSCHEL-like (*WUS*) and Transcription factor *TCP* in the South and distinct copies of *AP2/ERF* in the North and South.

**Figure 5.**
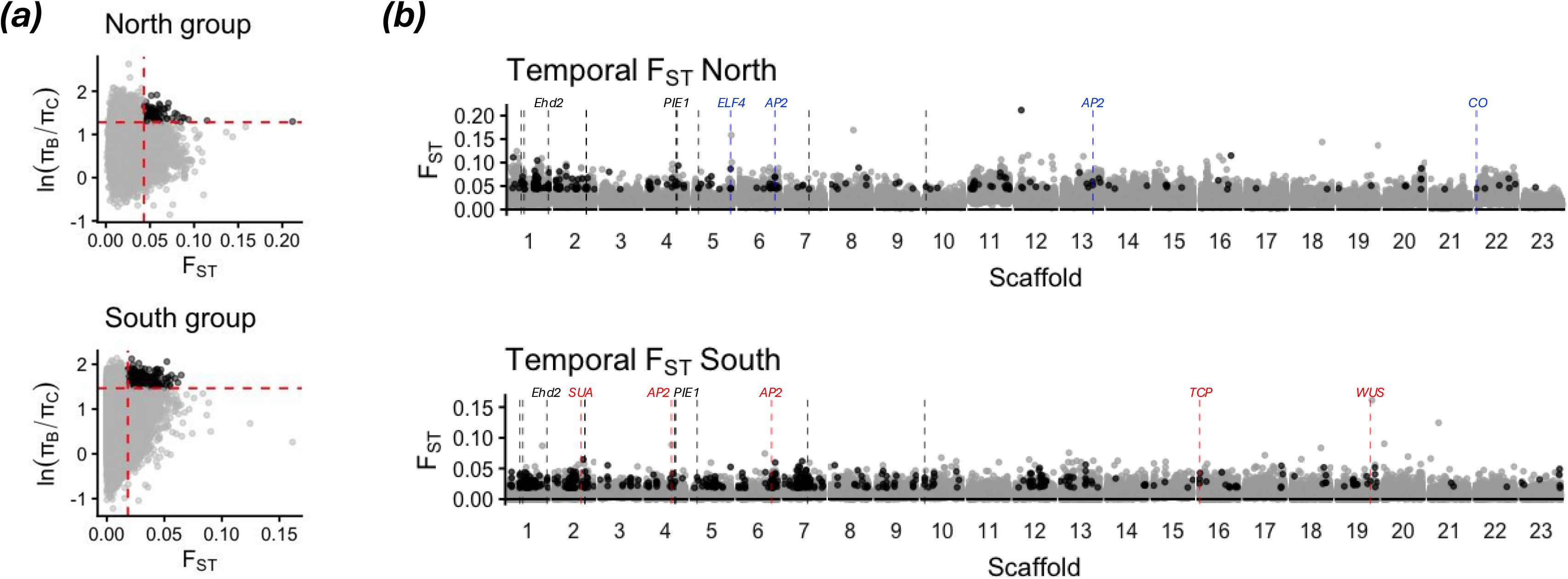
Putative selective sweeps in the introduced range. (a) Distributions of F_ST_ and nucleotide diversity ratios between samples collected before 1960 and contemporary samples for the North and South separately. Red dashed lines indicate the empirical 95^th^ percentile thresholds. (b) Genomic location of putative selective sweeps (black dots) and background windows (gray dots). Each dot represents a 10-kb window. Vertical dashed lines indicate the physical position of overlapping genes (black) or candidate genes for the North (blue) and South (red).

## Discussion

Biological invasions represent powerful natural experiments for studying evolution over ecological timescales. Until recently, however, our understanding of evolutionary processes occurring during biological invasions has been hindered by limited information on genotypes from the earliest stages of the invasion process. Here, we leveraged lcWGS of historical and contemporary samples of stiltgrass (*M. vimineum*) to reconstruct its invasion history in the eastern USA. Specifically, we employed a spatiotemporal approach to investigate the role of colonization history, admixture, and selection in invasion dynamics. Our analyses support multiple independent introductions to the USA involving an awnless form in the South and two awned forms in the North. Genetic diversity was highest in the earliest available collections, showing that substantial variation was already present early in the sampled invasion history. This pattern is compatible with short-lived demographic contractions and does not require an unobserved recovery of diversity. Admixture was overall low across space and time, and it is likely consequence of secondary contact due to rapid range expansion of separate introductions. Time series analysis of genetic structure revealed genetic continuity in both southern and northern regions and limited admixture between genetic groups, suggesting that habitat filtering may help maintain genetic groups (and phenotypic forms) in different regions of the introduced range. Finally, we found putative selective sweeps in the introduced range, with putative flowering-time genes in or near these genomic regions and limited but significant overlap between candidate genes in the northern and southern regions. Our study highlights the power of historical collections to uncover the evolutionary processes and mechanisms underlying successful biological invasions.

### Colonization history explains phenotypic distribution in the introduced range

Reconstructing invasion histories is critical for disentangling the evolutionary processes underlying successful invasions and for designing best management strategies (Estoup & Guillemaud, 2010; Bock *et al*., 2015; Cristescu, 2015; Kim *et al*., 2023). Here, we provide genetic evidence supporting multiple independent introductions; we identified three well-differentiated genetic groups in the introduced range, one in the southern region with awnless phenotype and two in the northern region, both with awned phenotypes. Demographic models indicate that genetic groups diverged well before the introduction to the USA, at least 27,000 years ago, supporting the hypothesis of genetic differentiation (and phenotypic variation) due to multiple independent introductions rather than purely post-introduction evolution in the introduced range (Barrett *et al*., 2022). This scenario of multiple introductions is also supported by a recent plastid haplotype analysis (Corbett, 2026), in which a single plastid haplotype has persisted in the southern USA while multiple, diverse haplotypes appeared over time in the northern USA, helping to explain the overall high genetic diversity found in the introduced range.

Stiltgrass was likely introduced in the USA as packing material for porcelain shipments from the native range in eastern Asia (Tu, 2000). Our time series analysis showed that all three genetic groups were present (and likely already established) in the USA before 1950s, around the time that plastics (first cellophane and then bubble wrap and Styrofoam) started replacing straw as packing material (Marty, 2020), which likely interrupted the main vector of stiltgrass introduction to the USA. The PCA and pairwise F_ST_ analyses suggest that the South Q2 group could have been introduced from Taiwan instead of Japan. Although Japanese porcelain could have been packed and exported from Taiwan (as Taiwan was under Japanese rule between the end of the First Sino-Japanese War and World War II, from 1895 to 1945; Dawley, 2008), to our knowledge, there are no historical records of porcelain shipments from Taiwan to the USA. In addition, Corbett (2026) did not find Taiwanese plastid haplotypes in samples from the southern region, which instead carried Japanese haplotypes in most cases. Moreover, Corbett (2026) found highly divergent Japanese and Chinese plastid haplotypes in the northern region, which is largely consistent with our *locator* results. Despite this, it is possible that the exact progenitor of introduced populations was not sampled in this study or even no longer exists in the native range. In addition, if stiltgrass was being moved by humans across continents, it is possible that different haplotypes were likewise moved across Asia, potentially obscuring patterns of native-range population structure.

### Low support for admixture as a driver of invasion

Our temporal approach revealed striking patterns of genetic diversity: the highest diversity was already present in the earliest available collections. Demographic modelling of contemporary samples inferred substantial population contractions near the period of introduction, but these results are not necessarily contradictory. The inferred contractions were brief (21–25 generations), and their effects on diversity depend on absolute effective population size as well as the magnitude of the decline. High early diversity could therefore reflect retention of founding variation, replenishment through subsequent introductions, or both. The earliest collections may also postdate initial establishment, because introduced populations can remain small and localized during a lag phase (Aikio *et al*., 2010). However, the sampled time series does not directly show a preceding loss and recovery of diversity. The low Tajima’s D values in the earliest time windows are consistent with subsequent expansion (Tajima, 1989), but do not by themselves demonstrate severe genetic bottlenecks. Together, the demographic and historical data show that this invasion was not characterized by uniformly low genetic diversity and illustrate the value of combining coalescent modelling with historical sampling.

Despite multiple introductions and clear secondary contact, we observed surprisingly low levels of admixture in the introduced range across time and space, i.e., most samples show high ancestry proportions from a single genetic group. Moreover, genetic divergence (F_ST_) between southern and northern regions increased over time, and geographically intermediate populations with short awns thought to be awnless/awned hybrids (Barrett *et al*., 2022) show little evidence of admixture. Thus, limited admixture may reflect restricted opportunities for interlineage mating, including selfing and reported differences in flowering time (Corbett, 2026), or habitat filtering associated with the strong covariance among genetic groups, traits, and geography (Barrett *et al*., 2022). Intrinsic reproductive barriers could also contribute, but high pairwise F_ST_ alone does not establish their presence. These mechanisms are not mutually exclusive, and we hypothesize that reproductive and ecological barriers both help maintain the observed genetic structure. Further experiments evaluating crosses between and within genetic groups in common gardens will help to disentangle the effect of these factors and to determine the potential benefits and limitations of admixture in this system. Taken together, the temporal and spatial patterns provide little support for extensive interlineage admixture having played a major role in establishment and range expansion; specifically, they do not support the strong prediction of the bottleneck–admixture hypothesis that admixture restored diversity and preceded range expansion in this system.

### Rapid evolution post-introduction

Rapid evolution is increasingly recognized as a major factor contributing to the success of invasive species (Prentis *et al*., 2008; Colautti & Barrett, 2013; Felker-Quinn *et al*., 2013; Hernández *et al*., 2019; Gamba *et al*., 2025; Nagy *et al*., 2026); however, evidence for *in situ* genetic changes occurring in the introduced range remains scarce (but see Bieker *et al*., 2022; Battlay *et al*., 2023, 2025). Here, we uncovered separate invasions in the southern and northern USA and identified putative selective sweeps in both regions, with limited but significant overlap between their candidate genes. Interestingly, among the overlapping genes, we found putative homologs of two flowering-time regulators: *Ehd2*, which promotes flowering through *Ehd1* in rice, and *PIE1*, which is required for *FLC* activation and floral repression in *Arabidopsis* (Noh & Amasino, 2003; Matsubara et al., 2008). Moreover, in the North group, we identified putative homologs of two additional genes (*ELF4* and *CO*) involved in flowering time regulation in response to photoperiod (Doyle *et al*., 2002), which is interesting because the North group is established at latitudes higher than most populations from the native range. Together with reported latitudinal clines in phenology (Novy *et al*., 2013; Corbett, 2026), our results suggest that selection on flowering time may have contributed to rapid adaptation and divergence of stiltgrass in the introduced range. Flowering time is among the most consistent traits showing rapid local adaptation in the introduced range, and likely facilitating range expansion (Colautti & Barrett, 2013; McGoey *et al*., 2020; Chen *et al*., 2023). Further studies using higher coverage and/or targeted sequencing of candidate genes coupled with common garden experiments are warranted to determine whether observed geographic clines in phenotypes and sequences facilitated range expansion and invasiveness of stiltgrass in the USA.

### Colonization history and selection jointly shape the invasion of stiltgrass

Although colonization history can explain phenotypic and genetic ancestry distributions, their strong association with geography, despite clear secondary contact, suggests that habitat filtering helps maintain phenotypic forms and genetic groups across different regions of the introduced range. Awns are hygroscopic structures that can move in response to changes in soil moisture content, contributing to seed dispersal and burial across grass species (Petersen & Kellogg, 2022). In stiltgrass, awns are thought to contribute to seed burial under freezing conditions during winter in northern latitudes; however, it is unclear whether the awnless phenotype is advantageous in the southern USA or whether the presence/absence of awns has fitness costs. An alternative explanation is that the presence/absence of awns is not adaptive itself, but it covaries strongly with adaptive traits like flowering time, plant architecture, or seed dormancy. Such covariance could be due to association with ancestry, i.e., genetic groups varying in presence/absence of awns also show differences in other traits (Barrett *et al*., 2022; Corbett, 2026), but also due to shared genetic mechanisms, as the presence/absence of awns is often regulated by conserved genes underlying plant development (Richardson *et al*., 2024). Further studies evaluating segregating populations for the awn phenotype under reciprocal common gardens are needed to disentangle the effects of environment, genetic ancestry, and phenotypic variation in shaping stiltgrass adaptation.

### Conclusions

Using historical and contemporary genomic data, we reconstructed the invasion history of *Microstegium vimineum* in the USA. Colonization history played a major role in shaping the present-day genetic and phenotypic patterns across the introduced range. The persistence of distinct genetic groups and limited admixture across space and time challenge a major role for admixture in invasion success. Substantial genetic variation was already present in the earliest available collections, and putative selective sweeps in or near flowering-time genes suggest rapid post-introduction evolution. Together, these findings highlight how historical collections reveal the distinct contributions of colonization history, standing variation and selection during invasion.

## Acknowledgements

This work was supported by the US National Science Foundation (award OIA-1920858), the West Virginia University (WVU) Department of Biology, the US Department of Agriculture Forest Service Northern Research Station, and the Science and Technology Projects of Xizang Autonomous Region, China (XZ202402ZD0005). For access to herbarium collections we thank Rehman, T. (BRIT), Isaac, B. (CM), Heberling, M. (CM), Freudenstein, J. (OS), Statler, A. (BH), Livshultz, T. (PH), Toner, M. (US), Boyle, L. (MO), Oliver, M. (TENN), Molano-Flores, B. (ILLS), Minnaert, J. (ILLS), Latvis, M. (UARK), Beck, J. (WSU), McKain, M. (UA) and Ford-Werntz, D. (WVA). We thank the following collaborators for providing field-collected material: Bowe, A., Hara, C., Crim, P., Fagan, J., Foster, B., Heberling, M., Isaac, B., Kennedy, A., Koenig, N., Kuebbing, S., Latvis, M., Matlack, G., McNeal, J., Molano-Flores, B., Moore, G., Mori, S., Puppo, P., Sato, S., Scott, G., Shimizu, T., Tanaka, K., Willard, K., and Wolf, P. For assistance with genomic sequencing, we thank Percifield, R., Primerano, D., Fan, J., Salisbury, T., and Sergent, B. We thank the WVU Genomics Core Facility for support provided to help make this publication possible and for CTSI grant U54 GM104942, which in turn provides financial support to the WVU Core Facility. We further acknowledge WV-INBRE (P20GM103434), a COBRE ACCORD grant (1P20GM121299), and a West Virginia Clinical and Translational Science Institute grant (2U54GM104942) in supporting the Marshall University Genomics Core (RRID:SCR_018885). The analyses were made possible by computational resources provided by the Digital Research Alliance of Canada.

## Competing interests

None to declare

## Author contributions

CFB and FH conceived the study, analyzed data, and led the writing of the manuscript. CDH, C-HC, BL, PL, PMP, KS, and W-BY provided samples and contributed to drafts of the manuscript. CWC and LEK conducted lab work, produced genomic sequence data, and contributed to drafts of the manuscript. FH, CFB, CWC, KS, and NJK analyzed data and helped draft the manuscript. All authors have reviewed and approved the final manuscript.

## Data availability

Analysis code is available at https://github.com/fernandoh33/Invasion-genomics-stiltgrass. Raw sequence reads are deposited in the NCBI Sequence Read Archive under BioProject PRJNA1479732. Sample metadata and analysis-specific inclusion criteria are provided in Table S1.

## Figure legends

**Figure S1.**
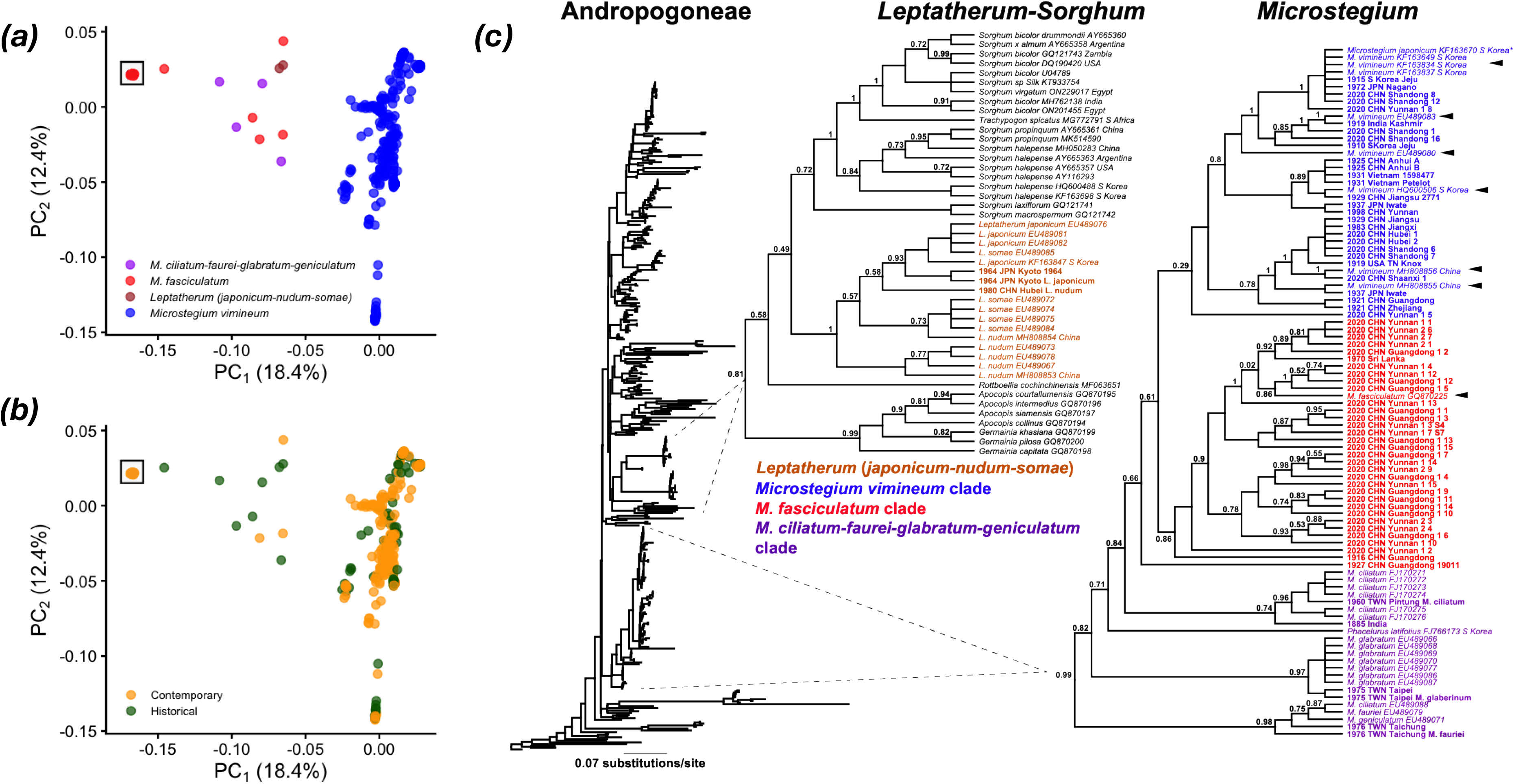
(a, b) Principal component analysis using all 472 samples colored by clade (a) or contemporary/historical category (b). The top-left square in both (a, b) highlights 29 contemporary samples collected in Guangdong and Yunnan provinces. (c) Maximum-likelihood phylogenetic analysis of 385 nuclear ITS accessions of grass tribe Andropogoneae. Phylogram of all taxa of Andropogoneae included (scale bar = 0.07 substitutions/site). Dotted lines correspond to clades in *Leptatherum-Sorghum* and *Microstegium*. Cladogram of the *Leptatherum-Sorghum* clade, with the *Leptatherum* clade highlighted in orange. Cladogram of *Microstegium*, with accessions in blue corresponding to *M. vimineum*, in red to *M. fasciculatum*, and in purple to *M. ciliatum, M. fauriei, M. glabratum*, and *M. geniculatum.* Numbers adjacent to branches are bootstrap frequencies. All accessions in bold were newly sequenced in the current study; names indicate the original year of collection (2019 onward = contemporary; before 2019 = historical herbarium samples). Black arrows in (c) correspond to GenBank reference accessions for the *M. vimineum* and *M. fasciculatum* clades.

**Figure S2.**
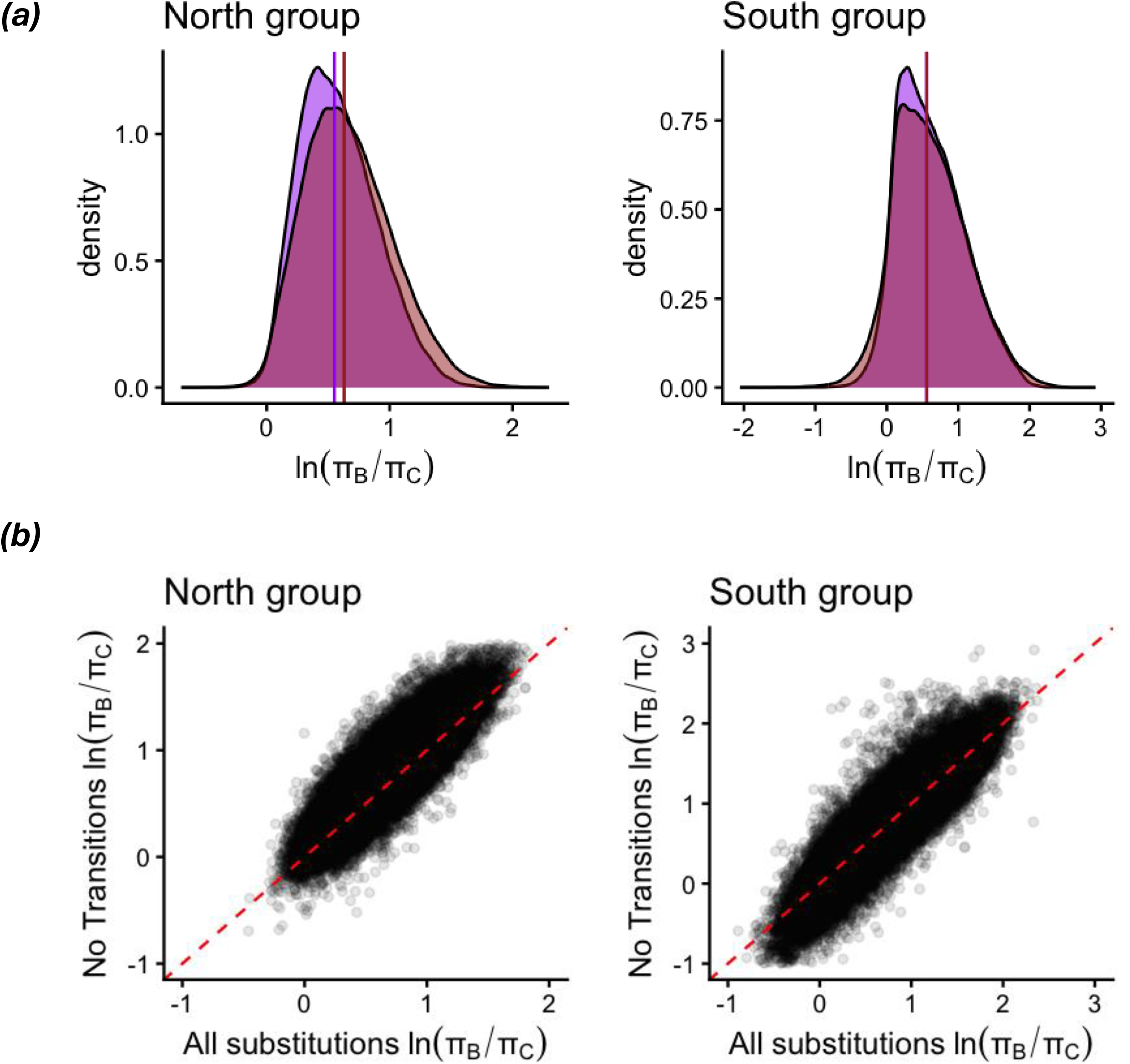
The reduction in nucleotide diversity between pre-1960 and contemporary samples persisted after transitions (A↔G and C↔T) were excluded to account for potential artifacts from ancient-DNA damage. (a) Genome-wide variation in the ratio between genetic diversity before 1960 and that in contemporary samples [ln(π_B_/ π_C_)] using all substitutions (purple) or excluding transitions (brown). Solid vertical lines indicate median values for each group. (b) Relationship between estimates of ln(π_B_/ π_C_) using all substitutions and excluding transitions. Each dot represents a 10-kb window with at least 1,000 sites in both groups; the red dashed line indicates the 1:1 relationship.

## Supplementary information

### Phylogenetic analysis

To explore the phylogenetic placement of accessions either known or suspected to belong to species other than *M. vimineum*, we conducted a phylogenetic analysis of the nuclear internal transcribed spacer (ITS). First, we downloaded all ITS accessions for the grass tribe Andropogoneae from NCBI GenBank (accessed 04 November 2025) and removed excessive duplicate members of the same species manually, resulting in 315 reference ITS sequences. We selected all newly sequenced historical samples from the native range (29) and all contemporary samples from China (31) for a total of 70 accessions. Paired-end sequences were trimmed with *fastp* (as above) and merged by overlap with bbmerge.sh of the BBTOOLS suite (Bushnell, 2014) using the ‘strict’ criterion for merging, trimming any remaining adapter overhangs (‘-tbo’ option) and discarding unmerged reads. Merged reads were then subsampled with seqkit to a maximum of 5 × 10^6^ reads per accession. Reads were imported into GENEIOUS v10 (https://www.geneious.com) and mapped 5× iteratively to the 5.8S region of *Microstegium japonicum* (GenBank number KF163847) to extend contigs to capture the flanking ITS1 and ITS2 regions (parameters: “Low sensitivity,” min overlap = 25, min overlap identity = 80%, max gaps per read = 10, max gap size = 3). Resulting consensus sequences were aligned with the GenBank references with the MAFFT (Katoh & Standley, 2013) plugin for GENEIOUS (gap open penalty = 3, offset value = 0.15). The resulting alignment ends were trimmed manually resulting in an alignment of 582 bp. Phylogenetic analysis was conducted using IQ-TREE 2 (Minh *et al*., 2020), with MODELFINDER (Kalyaanamoorthy *et al*., 2017) used to determine the best-fit model, and support assessed with 2,000 ultrafast bootstrap pseudoreplicates (UFBOOT2: Hoang *et al*., 2018).

The PCA separated samples into three main groups (Fig. **S1a,b**): one in the top left comprising 29 contemporary samples collected in China (Guangdong and Yunnan), one in the middle including three contemporary samples from Yunnan and eight historical samples collected in China, India, Japan, Sri Lanka, and Taiwan, and one in the right that includes all historical and contemporary samples from the introduced range, and all contemporary samples from the native range except the ones collected in Guangdong and Yunnan. Subsequent phylogenetic analysis shows that PCA groups largely correspond with sister-species clades (Table S1).

Phylogenetic analysis of the assembled nuclear ITS region from 70 accessions and 40 GenBank reference sequences from *Microstegium* (285 parsimony-informative characters, best-fit model = TIM2+R5) clearly differentiated a strongly supported *Leptatherum* clade (in orange; bootstrap percentage, or BP = 100), corresponding to *Leptatherum japonicum*, *Leptatherum nudum*, and *Leptatherum somae* (formerly placed in *Microstegium* but transferred to *Leptatherum*; Chen *et al*., 2009; 2012; Fig. **S1c**). Two of the historical accessions from Kyoto, Japan (one *L. japonicum* and one *L.* sp.) and one from Hubei, China (*L. nudum*) grouped within this clade, closely related to several accessions of *L. japonicum*. All other accessions grouped within a strongly supported clade corresponding to the currently circumscribed *Microstegium* (BS = 99). Two accessions from Taiwan grouped within a *M. ciliatum-fauriei-geniculatum* clade (in purple; BS = 98), and two others from Taiwan grouped with a *M. glabratum* clade (in purple; BS = 97). All other samples grouped in each of two other sister clades, excluding a 1927 accession from Guangdong, China, which grouped as sister to the aforementioned clades (collectively). The first clade (in red) included 30 contemporary accessions, all from Guangdong and Yunnan, China (BS = 86), within which a single reference accession of *M. fasciculatum* was nested (BS = 86). The clade also included historical accessions from Guangdong (1916, 1927) and Sri Lanka (1970). The other clade (in blue) corresponds to several reference accessions of *M. vimineum*, with several historical and contemporary accessions, likely representing a *M. vimineum* clade.

## Notes

### Competing Interest Statement

The authors have declared no competing interest.

## References

Aikio S, Duncan RP, Hulme PE. 2010. Lag-phases in alien plant invasions: separating the facts from the artefacts. Oikos 119: 370–378.

Alex Buerkle C, Gompert Z. 2013. Population genomics based on low coverage sequencing: How low should we go? Molecular Ecology 22: 3028–3035.

Baker HG. 1965. Characteristics and modes of origin of weeds. In The Genetics of Colonizing Species (pp. 147–169). Academic Press, New York.

Barker BS, Andonian K, Swope SM, Luster DG, Dlugosch KM. 2017. Population genomic analyses reveal a history of range expansion and trait evolution across the native and invaded range of yellow starthistle (*Centaurea solstitialis*). Molecular Ecology 26: 1131–1147.

Barker BS, Cocio JE, Anderson SR, Braasch JE, Cang FA, Gillette HD, Dlugosch KM. 2019. Potential limits to the benefits of admixture during biological invasion. Molecular Ecology 28: 100–113.

Barrett CF, Huebner CD, Bender ZA, Budinsky TA, Corbett CW, Latvis M, McKain MR, Motley M, Skibicki S V., Thixton HL, et al. 2022. Digitized collections elucidate invasion history and patterns of awn polymorphism in *Microstegium vimineum*. American Journal of Botany 109: 689–705.

Battey C, Ralph PL, Kern AD. 2020. Predicting geographic location from genetic variation with deep neural networks. eLife 9: e54507.

Battlay P, Hendrickson BT, Mendez-Reneau JI, Santangelo JS, Albano LJ, Wilson J, Caizergues AE, King N, Puentes A, Tudoran A, et al. 2025. Haploblocks contribute to parallel climate adaptation following global invasion of a cosmopolitan plant. Nature Ecology and Evolution 9: 1441–1455.

Barnett DW, Garrison EK, Quinlan AR, Strömberg MP, Marth GT. 2011. BamTools: a C++ API and toolkit for analyzing and managing BAM files. Bioinformatics 27: 1691–1692.

Battlay P, Wilson J, Bieker VC, Lee C, Prapas D, Petersen B, Craig S, van Boheemen L, Scalone R, de Silva NP, et al. 2023. Large haploblocks underlie rapid adaptation in the invasive weed *Ambrosia artemisiifolia*. Nature Communications 14: 1717.

Bieker VC, Battlay P, Petersen B, Sun X, Wilson J, Brealey JC, Bretagnolle F, Nurkowski K, Lee C, Barreiro FS, et al. 2022. Uncovering the genomic basis of an extraordinary plant invasion. Science Advances 8: eabo5115.

Blossey B, Nötzold R. 1995. Evolution of Increased Competitive Ability in Invasive Nonindigenous Plants: A Hypothesis. Journal of Ecology 83: 887–889.

Bock DG, Caseys C, Cousens RD, Hahn MA, Heredia SM, Hübner S, Turner KG, Whitney KD, Rieseberg LH. 2015. What we still don’t know about invasion genetics. Molecular Ecology 24: 2277–2297.

Chan YL, Anderson CNK, Hadly EA. 2006. Bayesian Estimation of the Timing and Severity of a Population Bottleneck from Ancient DNA. PLoS Genetics: e59.

Chen S, Zhou Y, Chen Y, Gu J. 2018. fastp: an ultra-fast all-in-one FASTQ preprocessor. Bioinformatics 34: i884–i890.

Chen X, Liu W, Zhang Y-Y, Zhang Y. 2023. Altered trait covariances between invasive and native ranges of a global plant invader. Functional Ecology 37: 1280–1290.

Cheplick G. 2010. Limits to local spatial spread in a highly invasive annual grass (*Microstegium vimineum*). Biological Invasions 12: 1759–1771.

Colautti RI, Barrett SCH. 2013. Rapid Adaptation to Climate Facilitates Range Expansion of an Invasive Plant. Science 342: 364–366.

Corbett CW. 2026. Genomic variation, adaptive evolution, and invasive and evolutionary history of *Microstegium vimineum*. Graduate Theses, Dissertations, and Problem Reports (ETD). https://researchrepository.wvu.edu/etd/13261.

Cristescu ME. 2015. Genetic reconstructions of invasion history. Molecular Ecology 24: 2212– 2225.

Danecek P, Bonfield JK, Liddle J, Marshall J, Ohan V, Pollard MO, Whitwham A, Keane T, McCarthy SA, Davies RM, Li H. 2021. Twelve years of SAMtools and BCFtools. GigaScience 10: giab008.

Dawley, E. 2008. Review of Taiwan under Japanese Colonial Rule, 1895–1945: History, Culture, Memory. Journal of Colonialism and Colonial History 9: 3.

Dlugosch KM, Anderson SR, Braasch J, Cang FA, Gillette HD. 2015. The devil is in the details: Genetic variation in introduced populations and its contributions to invasion. Molecular Ecology 24: 2095–2111.

Dlugosch KM, Parker IM. 2008. Founding events in species invasions: Genetic variation, adaptive evolution, and the role of multiple introductions. Molecular Ecology 17: 431–449.

Doyle JJ, Doyle JL. 1987. A rapid DNA isolation procedure for small quantities of fresh leaf tissue. Phytochemical Bulletin 19: 11–15.

Doyle M, Davis S, Bastow R, McWatters HG, Kozma-Bognár L, Nagy F, Millar AJ, Amasino RM. 2002. The *ELF4* gene controls circadian rhythms and flowering time in *Arabidopsis thaliana*. Nature 419: 74–77.

Early R, Bradley BA, Dukes JS, Lawler JJ, Olden JD, Blumenthal DM, Gonzalez P, Grosholz ED, Ibañez I, Miller LP, et al. 2016. Global threats from invasive alien species in the twenty-first century and national response capacities. Nature Communications 7: 12485.

Ellstrand NC, Schierenbeck KA. 2006. Hybridization as a stimulus for the evolution of invasiveness in plants? Euphytica 148: 35–46.

Enders M, Havemann F, Ruland F, Bernard-Verdier M, Catford JA, Gómez-Aparicio L, Haider S, Heger T, Kueffer C, Kühn I, et al. 2020. A conceptual map of invasion biology: Integrating hypotheses into a consensus network. Global Ecology and Biogeography 29: 978– 991.

Estoup A, Guillemaud T. 2010. Reconstructing routes of invasion using genetic data: Why, how and so what? Molecular Ecology 19: 4113–4130.

Estoup A, Ravigné V, Hufbauer R, Vitalis R, Gautier M, Facon B. 2016. Is There A Genetic Paradox of Biological Invasion? Annual Review of Ecology, Evolution, and Systematics 47: 51– 72.

Evanno G, Regnaut S, Goudet J. 2005. Detecting the number of clusters of individuals using the software STRUCTURE: a simulation study. Molecular Ecology 14: 2611–2620.

Excoffier L, Marchi N, Marques DA, Matthey-Doret R, Gouy A, Sousa VC. 2021. fastsimcoal2: demographic inference under complex evolutionary scenarios. Bioinformatics 37: 4882–4885.

Fairbrothers DE, Gray JR. 1972. *Microstegium vimineum* (Trin.) A. Camus (Gramineae) in the United States. Bulletin of the Torrey Botanical Club 99: 97.

Felker-Quinn E, Schweitzer JA, Bailey JK. 2013. Meta-analysis reveals evolution in invasive plant species but little support for Evolution of Increased Competitive Ability (EICA). Ecology and Evolution 3: 739–751.

Fick SE, Hijmans RJ. 2017. WorldClim 2: new 1km spatial resolution climate surfaces for global land areas. International Journal of Climatology 37: 4302–4315.

Fox EA, Wright AE, Fumagalli M, Vieira FG. 2019. ngsLD: evaluating linkage disequilibrium using genotype likelihoods. Bioinformatics 35: 3855–3856.

Fumagalli M. 2013. Assessing the effect of sequencing depth and sample size in population genetics inferences. PLoS ONE 8: e79667.

Gamba D, Vahsen ML, Maxwell TM, Pirtel N, Romero S, Van Ee JJ, Penn A, Das A, Ben-Zeev R, Baughman O, et al. 2025. Local adaptation to climate has facilitated the global invasion of cheatgrass. Nature Communications 16: 10203.

Hendrickson B, Patterson CM, King N, Doucet BI, Hernandez F, Kooyers NJ. 2026. Herbarium genomics reveal signatures of colonization history, lineage turnover, and adaptation during invasion. BioRxiv 10.64898/2026.03.30.715407.

Hernández F, Poverene M, Garayalde A, Presotto A. 2019. Re-establishment of latitudinal clines and local adaptation within the invaded area suggest rapid evolution of seed traits in Argentinean sunflower (*Helianthus annuus* L.). Biological Invasions 21: 2599–2612.

Hernández F, Vercellino RB, Todesco M, Bercovich N, Alvarez D, Brunet J, Presotto A, Rieseberg LH. 2024. Admixture With Cultivated Sunflower Likely Facilitated Establishment and Spread of Wild Sunflower (*Helianthus annuus*) in Argentina. Molecular Ecology 33: e17560.

Hodgins KA, Battlay P, Bock DG. 2025. The genomic secrets of invasive plants. New Phytologist. 245: 1846–1863.

Hovick SM, Whitney KD. 2014. Hybridisation is associated with increased fecundity and size in invasive taxa: Meta-analytic support for the hybridisation-invasion hypothesis. Ecology Letters 17: 1464–1477.

Innes SG, Santangelo JS, Kooyers NJ, Olsen KM, Johnson MTJ. 2022. Evolution in response to climate in the native and introduced ranges of a globally distributed plant. Evolution 76: 1495–1511.

Jónsson H, Ginolhac A, Schubert M, Johnson PLF, Orlando L. 2013. mapDamage2.0: fast approximate Bayesian estimates of ancient DNA damage parameters. Bioinformatics 29: 1682– 1684.

Keller SR, Fields PD, Berardi AE, Taylor DR. 2014. Recent admixture generates heterozygosity-fitness correlations during the range expansion of an invading species. Journal of Evolutionary Biology 27: 616–627.

Keller SR, Taylor DR. 2008. History, chance and adaptation during biological invasion: Separating stochastic phenotypic evolution from response to selection. Ecology Letters 11: 852– 866.

Kim AS, Kreiner JM, Hernández F, Bock DG, Hodgins KA, Rieseberg LH. 2023. Temporal collections to study invasion biology. Molecular Ecology 32: 6729–6742.

Kopelman NM, Mayzel J, Jakobsson M, Rosenberg NA, Mayrose I. 2015. Clumpak: a program for identifying clustering modes and packaging population structure inferences across K. Molecular Ecology Resources 15: 1179–1191.

Korneliussen TS, Albrechtsen A, Nielsen R. 2014. ANGSD: Analysis of Next Generation Sequencing Data. BMC Bioinformatics 15: 356.

Kreiner JM, Latorre SM, Burbano HA, Stinchcombe JR, Otto SP, Weigel D, Wright SI. 2022. Rapid weed adaptation and range expansion in response to agriculture over the past two centuries. Science 378:1079–1085.

Lang PLM, Willems FM, Scheepens JF, Burbano HA, Bossdorf O. 2019. Using herbaria to study global environmental change. New Phytologist 221: 110–122.

Li H, Durbin R. 2009. Fast and accurate short read alignment with Burrows-Wheeler transform. Bioinformatics 25: 1754–1760.

Liu C, Wolter C, Xian W, Jeschke JM. 2020. Most invasive species largely conserve their climatic niche. Proceedings of the National Academy of Sciences 117: 23643–23651.

Lockwood JL, Cassey P, Blackburn T. 2005. The role of propagule pressure in explaining species invasions. Trends in Ecology & Evolution 20: 223–228.

Lou RN, Jacobs A, Wilder A, Therkildsen NO. 2021. A beginner’s guide to low-coverage whole genome sequencing for population genomics. Molecular Ecology 30: 5966–5993.

Lou RN, Therkildsen NO. 2022. Batch effects in population genomic studies with low-coverage whole genome sequencing data: Causes, detection and mitigation. Molecular Ecology Resources 22: 1678–1692.

Lowry DB, Hoban S, Kelley J, Lotterhos KE, Reed LK, Antolin MCF, Torfer A. 2017. Breaking RAD: an evaluation of the utility of restriction site-associated DNA sequencing for genome scans of adaptation. Molecular Ecology Resources 17: 142–152.

Marty N. 2020. The True Revolution of 1968: Mineral Water Trade and the Early Proliferation of Plastic, 1960s–1970s. Business History Review 94: 483–505.

McGoey BV, Hodgins KA, Stinchcombe JR. 2020. Parallel flowering time clines in native and introduced ragweed populations are likely due to adaptation. Ecology and Evolution 10: 4595– 4608.

Matsubara K, Yamanouchi U, Wang ZX, Minobe Y, Izawa T, Yano M. 2008. *Ehd2*, a rice ortholog of the maize *INDETERMINATE1* gene, promotes flowering by up-regulating *Ehd1*. Plant Physiology 148: 1425–35.

Meisner J, Albrechtsen A. 2018. Inferring population structure and admixture proportions in low-depth NGS data. Genetics 210: 719–731.

Mirchandani CD, Shultz AJ, Thomas GWC, Smith SJ, Baylis M, Arnold B, Corbett-Detig R, Enbody E, Sackton TB. 2024. A Fast, Reproducible, High-Throughput Variant Calling Workflow for Population Genomics. Molecular Biology and Evolution 41: msad270.

Muhlfeld CC, Kovach RP, Jones LA, Al-Chokhachy R, Boyer MC, Leary RF, Lowe WH, Luikart G, Allendorf FW. 2014. Invasive hybridization in a threatened species is accelerated by climate change. Nature Climate Change 4: 620–624.

Nagy DU, Callaway RM, Luke Flory S, Barratt CD, Durka W, Lekberg Y, Lucas MS, Michalski SG, Onstein RE, Pal RW, et al. 2026. Rapid and repeated evolution of increased competitive ability in a global invader. Nature Communications 17: 9930.

Nei M, Maruyama T, Chakraborty R. 1975. The Bottleneck effect and genetic variability in populations. Evolution 29: 1–10.

Noh YS, Amasino RM. 2003. PIE1, an ISWI family gene, is required for FLC activation and floral repression in Arabidopsis. The Plant Cell 15:1671–1682.

North HL, McGaughran A, Jiggins CD. 2021. Insights into invasive species from whole-genome resequencing. Molecular Ecology 30: 6289–6308.

Novy A, Flory SL, Hartman JM. 2013. Evidence for rapid evolution of phenology in an invasive grass. Journal of Evolutionary Biology 26: 443–450.

Okonechnikov K, Conesa A, García-Alcalde F. 2016. Qualimap 2: advanced multi-sample quality control for high-throughput sequencing data. Bioinformatics 32: 292–294.

Pascual-Díaz JP, Besolí N, López-Pujol J, Nualart N, Pérez-Lorenzo I, Shtein R, Valenzuela L, Garcia S, Vitales D. 2025. The Winner Takes It All: a single genotype of *Kalanchoe* × houghtonii is a global invader. Annals of Botany 136: 179–198.

Petersen KB, Kellogg EA. 2022. Diverse ecological functions and the convergent evolution of grass awns. American Journal of Botany 109: 1331–1345.

Prentis P, Wilson J, Dormontt E, Richardson DM, Lowe AJ. 2008. Adaptive evolution in invasive species. Trends in Plant Science 13: 288–294.

Presotto A, Hernández F, Vercellino RB, Kruger RD, Fontana ML, Ureta MS, Crepy M, Auge G, Caicedo A. 2024. Introgression from local cultivars is a driver of agricultural adaptation in Argentinian weedy rice. Molecular Ecology 33: e17368.

Presotto A, Hernández F, Mercer KL. 2019. Phenotypic selection under two contrasting environments in wild sunflower and its crop–wild hybrid. Evolutionary Applications 12: 1703– 1717.

Pyšek P, Richardson DM. 2010. Invasive Species, Environmental Change and Management, and Health. Annual Review of Environment and Resources 35: 25–55.

Qiu J, Jia L, Wu D, Weng X, Chen L, Sun J, Chen M, Mao L, Jiang B, Ye C, et al. 2020. Diverse genetic mechanisms underlie worldwide convergent rice feralization. Genome Biology 21: 70.

Quinlan AR, Hall IM. 2010. BEDTools: A flexible suite of utilities for comparing genomic features. Bioinformatics 26: 841–842.

Ramachandran D, Huebner CD, Daly M, Haimovitz J, Swale T, Barrett CF. 2021. Chromosome Level Genome Assembly and Annotation of Highly Invasive Japanese Stiltgrass (*Microstegium vimineum*). Genome Biology and Evolution 13: evab238.

Rawlence NJ, Kardamaki A, Easton LJ, Tennyson AJD, Scofield RP, Waters JM. 2017. Ancient DNA and morphometric analysis reveal extinction and replacement of New Zealand’s unique black swans. Proceedings of the Royal Society B: Biological Sciences 284: 20170876.

Redwood ME, Matlack GR, Huebner CD. 2018. Seed Longevity and Dormancy State Suggest Management Strategies for Garlic Mustard (*Alliaria petiolata*) and Japanese Stiltgrass (*Microstegium vimineum*) in Deciduous Forest Sites. Weed Science 66: 190–198.

Rejmánek M, Richardson DM. 1996. What Attributes Make Some Plant Species More Invasive? Ecology 77: 1655–1661.

Richardson A, Jones H, Bartlett M. 2024. Grass awns: Morphological diversity arising from developmental constraint. Current Opinion in Plant Biology 82: 102663.

Rizzieri YC, Limpanasittichai P, Hernández F, Mendez-Reneau JI, Holt SD, Sutherland B, Mandáková T, Beck JB, Kuo L-Y, Kooyers NJ, et al. 2026. The dynamic genomes of *Salvinia* reshape our understanding of fern chromosome evolution. Proceedings of the National Academy of Sciences 123: e2602084123.

Sherpa S, Després L. 2021. The evolutionary dynamics of biological invasions: A multi-approach perspective. Evolutionary Applications 14: 1463–1484.

Skotte L, Korneliussen TS, Albrechtsen A. 2013. Estimating individual admixture proportions from next generation sequencing data. Genetics 195: 693–702.

Tajima, F. 1989. Statistical method for testing the neutral mutation hypothesis by DNA polymorphism. Genetics, 123: 585–595.

Tu, M. 2000. Element stewardship abstract for *Microstegium vimineum*. The Nature Conservancy. Website: https://www.invasive.org/gist/esadocs/documnts/micrvim.pdf.

Turner KG, Hufbauer RA, Rieseberg LH. 2013. Rapid evolution of an invasive weed. New Phytologist 202: 309–321.

Vahsen ML, Shea K, Hovis CL, Teller BJ, Hufbauer RA. 2018. Prior adaptation, diversity, and introduction frequency mediate the positive relationship between propagule pressure and the initial success of founding populations. Biological Invasions 20: 2451–2459.

van Boheemen LA, Lombaert E, Nurkowski KA, Gauffre B, Rieseberg LH, Hodgins KA. 2017. Multiple introductions, admixture and bridgehead invasion characterize the introduction history of *Ambrosia artemisiifolia* in Europe and Australia. Molecular Ecology 26: 5421–5434.

van Kleunen M, Röckle M, Stift M. 2015. Admixture between native and invasive populations may increase invasiveness of *Mimulus guttatus*. Proceedings of the Royal Society B: Biological Sciences 282: 20151487.

Vercellino RB, Hernández F, Presotto A. 2023. The role of intraspecific crop–weed hybridization in the evolution of weediness and invasiveness: Cultivated and weedy radish (*Raphanus sativus*) as a case study. American Journal of Botany 110: e16217.

Verhoeven KJF, MacEl M, Wolfe LM, Biere A. 2011. Population admixture, biological invasions and the balance between local adaptation and inbreeding depression. Proceedings of the Royal Society B: Biological Sciences 278: 2–8.

Wales N, Akman M, Watson RHB, Sánchez Barreiro F, Smith BD, Gremillion KJ, Gilbert MTP, Blackman BK. 2019. Ancient DNA reveals the timing and persistence of organellar genetic bottlenecks over 3,000 years of sunflower domestication and improvement. Evolutionary Applications 12: 38–53.

Weiher E, Keddy PA. 1995. Assembly Rules, Null Models, and Trait Dispersion: New Questions from Old Patterns. Oikos 74: 159.

Wickham H. 2016. ggplot2: Elegant Graphics for Data Analysis. Springer-Verlag New York. ISBN 978-3-319-24277-4. https://ggplot2.tidyverse.org.

Ziska LH, Tomecek MB, Valerio M, Thompson JP. 2015. Evidence for recent evolution in an invasive species, *Microstegium vimineum*, Japanese stiltgrass. Weed Research 55: 260–267.

## References

Bushnell, B. 2014. BBMap: A Fast, Accurate, Splice-Aware Aligner. GitHub: https://github.com/bbushnell/BBTools.

Chen C-H, Veldkamp JF, Kuoh C-S, Tsai C-C, Chiang Y-C. 2009. Segregation of *Leptatherum* from *Microstegium* (Andropogoneae, Poaceae) confirmed by Internal Transcribed Spacer DNA sequences. Blumea: Biodiversity, Evolution and Biogeography of Plants 54: 175–180.

Chen C-H, Veldkamp JF, Kuoh C-S. 2012. Taxonomic revision of *Microstegium* s.str. (Andropogoneae, Poaceae). Blumea: Biodiversity, Evolution and Biogeography of Plants 57: 160–189.

Hoang DT, Chernomor O, von Haeseler A, Minh BQ, Vinh LS. 2018. UFBoot2: Improving the Ultrafast Bootstrap Approximation, Molecular Biology and Evolution 35: 518–522.

Kalyaanamoorthy S, Minh BQ, Wong TKF, von Haeseler A, Jermiin LS. 2017. ModelFinder: fast model selection for accurate phylogenetic estimates. Nature Methods 14: 587–589.

Katoh K, Standley DM. 2013. MAFFT multiple sequence alignment software version 7: improvements in performance and usability. Molecular Biology and Evolution 30: 772–80.

Minh BQ, Schmidt HA, Chernomor O, Schrempf D, Woodhams MD, von Haeseler A, Lanfear R. 2020. IQ-TREE 2: New Models and Efficient Methods for Phylogenetic Inference in the Genomic Era. Molecular Biology and Evolution 37: 1530–1534.

